# Precise implantation of super-paramagnetic iron oxide (SPIO) nanoparticles in hippocampus with external rotating magnetic field (RMF) produces rapid antidepressent effects in mice

**DOI:** 10.1101/665315

**Authors:** Yao Chi, Cai Wenwen, Xia Mengqin, Dai Jingyi, JF Sun Sr.

## Abstract

**Background:** TMS is an effective anti-depression method commonly used in clinical practice, but it also faces the problems of low spatial resolution, treatment parameters to be optimized and limitation in mechanism research.

**Objective:** To develop a precise magnetic stimulation anti-depression method for the scientific research of magnetic stimulation, especially the mechanism research in animal experiments.

**Methods:** The cytotoxicity test was conducted in advance to ensure the security of intervention from SPIO nanoparticles and RMF. In animal experiments, 300nl SPIO solution was injected into the right hippocampus of the CUMS model mice, and then treated experimental group mice with rotating magnetic field for five days. The Sucrose Preference Test (SPT), the Forced Swim Test (FST) and BDNF expression levels were used to evaluate the antidepressant effect of this method.

**Results:** No significant decrease of cell viability was observed when the iron concentration is between 52.2µg/ml and 208.8µg/ml. And the application of RMF with a certain frequency was considered to be safe in the cytotoxicity test. When treated with SPIO+RMF, the sucrose preference of SPIO+RMF group mice increased markedly (n=9, p<0.01 vs. CUMS), the FST immobile time reduced (n=8, P<0.05 vs. CUMS) and the BDNF level in the hippocampus was significantly up-regulated (n=5, P<0.01 vs. CUMS). However, merely SPIO intervention failed to be effective.

**Conclusions:** when intervened with external rotating magnetic field, the SPIO nanoparticles injected into the right hippocampus could produce rapid antidepressent effects in mice.

## Introduction

As a prevalent and recurrent psychiatric disorder, major depressive disorder (MDD) evaluates risk of disability and early mortality [1,2]. According to WHO, it moved into the top 3 causes of the global burden of disease in 2004 and was projected to be the first by 2030 [3]. However, the first-line antidepressants like selective serotonin reuptake inhibitors (SSRIs) take moderate efficacy in several weeks and high placebo responses are included, which clinically contributes much to the low cure rate of MDD [4,5]. Therefore, there is an urgent need to develop and optimize novel treatments for depression.

In newer anti-depressive strategies, repetitive transcranial magnetic stimulation (rTMS) is one of efficient physical therapies as a non-invasive nerve stimulation technology [6]. It works with a coil of wire near the scalp which delivers repetitive pulse magnetic field to produce induced current, changing the membrane potential in nerve cells and causing a series of physiological and biochemical reaction [7]. From 2008 to 2015, four different rTMS devices for MDD patients had been declared by the United States Food and Drug Administration (FDA). However, the promising technique seems to trap into marsh when comes to cellular, molecular and neural circuit mechanisms due to the dilemma of magnetic field focusing accuracy. Factually, the smallest one of commercial human coils even stimulates the whole brain of rodent and rodent-specific TMS develops sluggishly for the conflict between size and intensity [8]. The stimulus intensity has to compromise if focal stimulation is delivered [9]. To target the pathophysiology linked brain regions, navigated TMS based on medical imaging technique [10] and deep transcranial magnetic stimulation (DTMS) equipped with the H-coil system [11] are developed, especially after Uwe Herwig et al. found the standard procedure of TMS is not precise anatomically [12]. Also, studies showed evidence that progresses were made in region targeted treatments [13,14]. However, general neuronavigation systems are limited for the heavy economic burden [15] and DTMS achieves deeper stimulation at the cost of influencing wider brain volumes [12]. Both the eager desire for mechanism and the predicament faced by TMS inspire us to turn to super-paramagnetic iron oxide (SPIO) nanoparticles.

Performance of SPIO nanoparticles in biomedical applications mounts [16, 17, 18, 19, 20]. Due to the superparamagnetic character, SPIO nanoparticles are widely used as contrast-enhancing agents for magnetic resonance imaging (MRI) [21, 22]. Moreover, the US FDA has approved several SPIO nanoparticles, (e.g., Feridex) [23]. Consequently, can the nanoparticle with magnetic properties be employed as accurate positioning magnetic stimulations under an applied rotating magnetic field(RMF) when injected into brain tissue whether cerebral cortex or deeper regions?

It should be verified whether the potential anti-depressive effect exists or not. As to SPIO-enhanced magnetic stimulation, cytotoxicity is tested in advance. Then we designed a protocol to investigate the effects on mice with depression-like behavior. Hippocampus, a region repeatedly implicated in the pathophysiology and depression progression [24], was chosen as our targeted brain region. It shows antidepressive-like effects in CUMS rodent model after a 5-day treatment. The approach developed in this paper may throw new light in quickly elevating mood for depression and provide a new perspective to study the mechanism of magnetic stimulation therapy.

## Materials and Methods

### Mice

In our study, we used 6- to 8-week-old male C57BL/6J mice (18-24g), purchased from Sino-British SIPPR/BK Lab Animal Ltd (Shanghai, China). All mice were group housed under standard conditions (12/12 hour light/dark cycle; light on from 07:00 AM to 7:00 PM; 23 ± 1°C ambient temperature; 55 ± 2% relative humidity) for 1 week with free access to food and water. All animal experiments were carried out in accordance with the National Institutes of Health (NIH) Guide for the Care and Use of Laboratory Animals and approved by Animal Study Committee of Southeast University, China.

### Equipment

The materials we used were rotating magnetic field and superparamagnetic iron oxide nanoparticles. The SPIO nanoparticle here we used held a γ-Fe2O3 core about 6-8nm and coated with polyglucose sorbitol carboxymethyether about 14nm. The aqueous dispersion was prepared according to the classic chemical co-precipitation method using ferrous chloride hexahydrate and ferrous chloride tetra-hydrate (SigmaeAldrich, MO, US). For physicochemical properties, the hydrodynamic size is 34 nm and the polydispersity index is 0.2. The saturation magnetization of SPIO nanoparticle is 77emu/g. The concentration of Fe element was 29.34mg/mL. It is supposed that it can take advantage of it’s superparamagnetic properties and then enhance the local magnetic field intensity by externally imposing a mild magnetic field when injected into hippocampus. In short, refer to other studies in our laboratory, we separately precisely injected 300nl SPIO solution into the right and bilateral hippocampus of mice using a stereotaxic apparatus. Then we used 10Hz RMF to intervene the corresponding group with magnetic head 5mm away from the mouse brain. We treated the mice twice a day with an eight-hour interval. Each treatment lasts for 5 minutes and the total course is 5 days.

### Cytotoxicity test

HT22 murine hippocampal neuronal cells (from Berke Biology Firm in Nanjing, Jiangsu Province, China) were maintained in H-DMEM (Dulbecco’s modified Eagle’s medium) supplemented with 10% FBS (GIBCO 10270-106) and antibiotics (Penicillin/streptomycin solution), and incubated at 37°C under 5% CO2. Cells were subcultured before every test for 7 days and seeded in 96-well plates at a density of 5000 cells per well. All SPIO nanoparticles were attenuated evenly by above medium while cells in control groups were treated with vehicle only.

Cell viability was measured by Cell Counting Kit-8 (CCK-8) (EnoGene Biology, Nanjing), according to the manufacturer’s instruction. Cells were divided into four groups by deferent magnetic field frequency, 0Hz, 2.5Hz, 5Hz and 10Hz, and Fe concentrations in per group varied from 26.1ug/ml to 417.5ug/ml. After an intervention for a 5 days, 10 ml of CCK-8 solution replaced the original medium to incubate for 2 h at 37°C. Finally, OD value that reflects the cell viability was read at 450nm with an enzyme-linked immunesorbent assay (ELISA) reader (KAYTO).

### CUMS modeling

CUMS is a classic and well-validated model of depression [25, 26]. Therefore we selected this animal model to verify the effect of SPIO+RMF on depression-like behavior. Before the start of modeling, we had made the mice adapted to the standard conditions and to drink in two tubes (two tubes of 1% fresh sucrose solution in the first two days, one tube of 1% fresh sucrose solution and one tube of fresh water in the next two days, two tubes of fresh water in the last three days, exchange the position of two tubes every day) for one week. Then we measured the SPT and FST baseline and excluded mice whose sucrose preference (sucrose/ total consumption*100%) were lower than 75% [27].

The process of model building lasted for 5 weeks. Mice were exposed to a variety of unpredictable and mild stressors, including restraint (2h), cold water swimming (4 °C, 5 min), warm water swimming (45 °C, 5 min), cage shaking (5min), tail pinch (1min), tail suspension (6min), food and/or water deprivation (12h), wet bedding (12h), overnight illumination (12h), tilted cage (10h), empty cage (12h) and stroboscopic illumination (12h) (see Table 1 for the schedule).

**Table 1.** Arrangements for daily stressors during the CUMS modeling. A:restraint (2h), B: cold water swimming (4 °C, 5 min), C: warm water swimming (45 °C,5 min), D: cage shaking (5min), E: tail pinch (1min), F: tail suspension (6min), G: water deprivation (12h), H: food and water deprivation (12h), I: wet bedding (12h), J: overnight illumination (12h), K: tilted cage (10h), L: empty cage (12h),M: stroboscopic illumination (12h).

| Days<br>weeks | 1 | 2 | 3 | 4 | 5 | 6 | 7 |
| --- | --- | --- | --- | --- | --- | --- | --- |
| 1 | AG | CJ | DK | EI | FL | DM | AH |
| 2 | KF | EB | DJ | MI | AL | EH | FJ |
| 3 | AK | DJ | CG | EM | FI | DL | AH |
| 4 | EB | DK | MG | AI | FL | DH | EJ |
| 5 | DK | EC | AI | LH | FM | DK | AH |

### Stereotactic microinjection and magnetic treatment

All successful model mice were assigned to three groups, including CUMS group, SPIO group and SPIO+RMF group. The sample size for each group was 8 or 9. For SPIO group and SPIO+RMF group mice, they were narcotized with 1% pentobarbital sodium (35mg-40mg/kg of body weight, i. p.) and were fixed in the stereotaxic apparatus next. They were injected 300nl SPIO at the rate of 200nl/min into the right hippocampus. According to The Mouse Brain in Stereotaxic Coordinates [28], the stereotaxic coordinates were: −2.3mm at the anterior/posterior axis, −1.6mm at the lateral/medial axis and −1.8mm at the dorsal/ventral axis. The whole process was slow and gentle to prevent back flows and bleeding [29]. The mice in SPIO+RMF group received RMF in the next 5 days as mentioned above while the SPIO group mice didn’t. To rule out spontaneous recovery, all groups continued to be exposed to stressors in the procedure of injection and treatment. (See Figure 1 for the procedure of animal experiments.)

**Figure 1.**
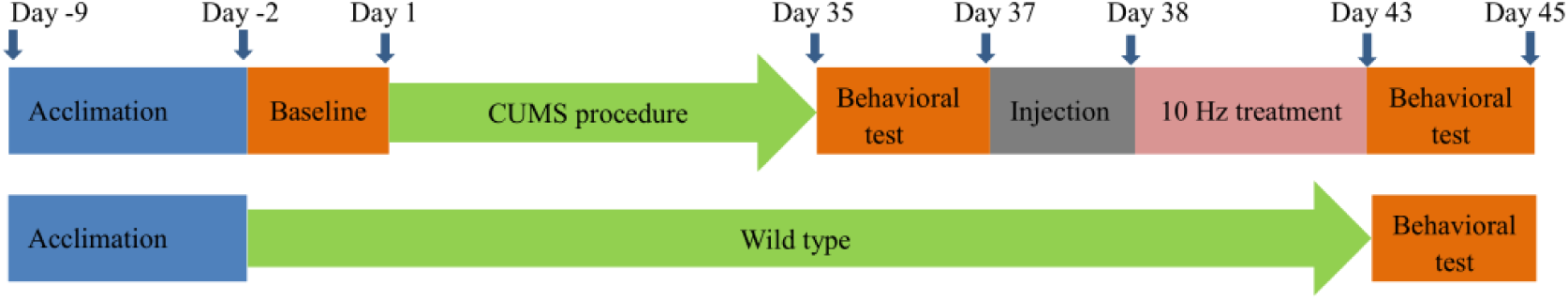
Experimental procedures for the CUMS modeling treatment and behavioral assessment.

### SPT

As is known to us, anhedonia is a typical symptom of depression [27]. In our study, we used SPT to quantify the degree of anhedonia. Considering that the absolute intake value of sucrose solution can fluctuate due to the weight variance of mice, we chosed the preference ratio of sucrose as the indicative parameter of the SPT. All the mice had experienced the process of sucrose solution adaptation in advance. The mice were deprived of food and water for 24h before the test. Then a 2h test of sucrose preference was performed (One tube in fresh water and the other in 1% fresh sucrose solution. Exchange the position of two tubes in the middle to prevent position preference.). Two hours later, we removed the tubes and measured the consumption of water and sucrose solution, calculated sucrose preference value according the following formula: sucrose preference (%) = [sucrose intake (g) / (sucrose intake + water intake) (g)] × 100%.

### FST

The forced swim test is a rodent behavioral test used for evaluation of antidepressant drugs, antidepressant efficacy of new compounds, and experimental manipulations that are aimed at rendering or preventing depressive-like states [30]. Mice were placed individually in a vertical glass cylinder (height, 25 cm; diameter, 12 cm) containing water about 15 cm deep at 21±1°C and forced to swim. We defined the first 2 minutes as the mouse adaptation time and the immobility time was measured during the last 4 minutes. The behaviors of each mouse were videotaped analyzed with ANY-MAZE behavior monitoring system.

### Western Blot

BDNF is a member of the structurally and functionally homologous neurotrophin family. It has been reported that decreased level of BDNF in hippocampus may alter the function and structure of hippocampal neurons, inducing depressive behaviors in mice [31]. Clinical studies previous also showed that BDNF level of patients with bipolar disorder and major depression decreased, which may recover after anti-depressive treatment [32]. Moreover, numerous experimental compounds and drugs are suggested to improve depressive-like behaviors by altering the protein expression, or the signaling pathway of BDNF at a deeper level [33, 34, 35]. Corresponding to compounds and drugs, BDNF was reported to be correlated with the antidepressive effects of rTMS [36]. A review calimed that BDNF is the most promising predictor of MDD victims’ respondence to TMS[37]. Therefore, BDNF was chosen as the molecular index for the study. The mice were immediately sacrificed in deep anesthesia with pentobarbital after the final behavioral test. The whole brain was collected and perfused with cold phosphate-buffered saline (PBS). We then extracted the right hippocampus samples and quantified BDNF protein levels. Protein quantification was analysis on ImageJ and the specific protein expression levels were normalized to the levels of GAPDH on the same PVDF membrane.

### Statistical analysis

All datas were expressed as means ± SE and were analyzed using Student’s t –test or One-way ANOVA followed by Bonferroni’s multiple comparisons post hoc test. Differences with p <0.05 between experimental groups at each point were considered statistically significant. Figures were obtained by the Statistical Analysis System (GraphPad Prism 7, GraphPad Software Inc., USA).

## Results

### 1. SPIO had little toxicity on cell viability within a certain range in vitro

#### Cytotoxicity test

A 5-day cytotoxicity effect of SPIO nanoparticles was firstly evaluated by CCK-8 approach utilizing HT22, the immortalized mouse hippocampal cell line, which is a candidate model to determinate the potential oxidative toxicity of SPIO [38, 39, 40, 41, 42]. A series of concentrations were set, they are 1/4, 1/8, 1/16, 1/32 and 1/64 of what we injected into hippocampus. Cell viability was also determined under the RMF at 2.5Hz, 5Hz and 10Hz. Results showed a concentration-dependent cytotoxicity. In the absence of RMF, cells incubated with different concentrations of SPIO solution possessed a viability of 54%, 78%, 84%, 88% and 77% compared with control group (Figure 2 A). As can be seen from figure 2(B,C,D), although the concentration-dependent toxicity still existed after the application of RMF, there were optimal internals where cell viability was similar to control groups or even higher (2.5hz, p < 0.05 vs. the control group). Interestingly, the concentration condition with the highest viability decreased as the increase of magnetic field frequency, suggesting that there was a corresponding relationship between RMF and SPIO concentration.

**Figure 2.**
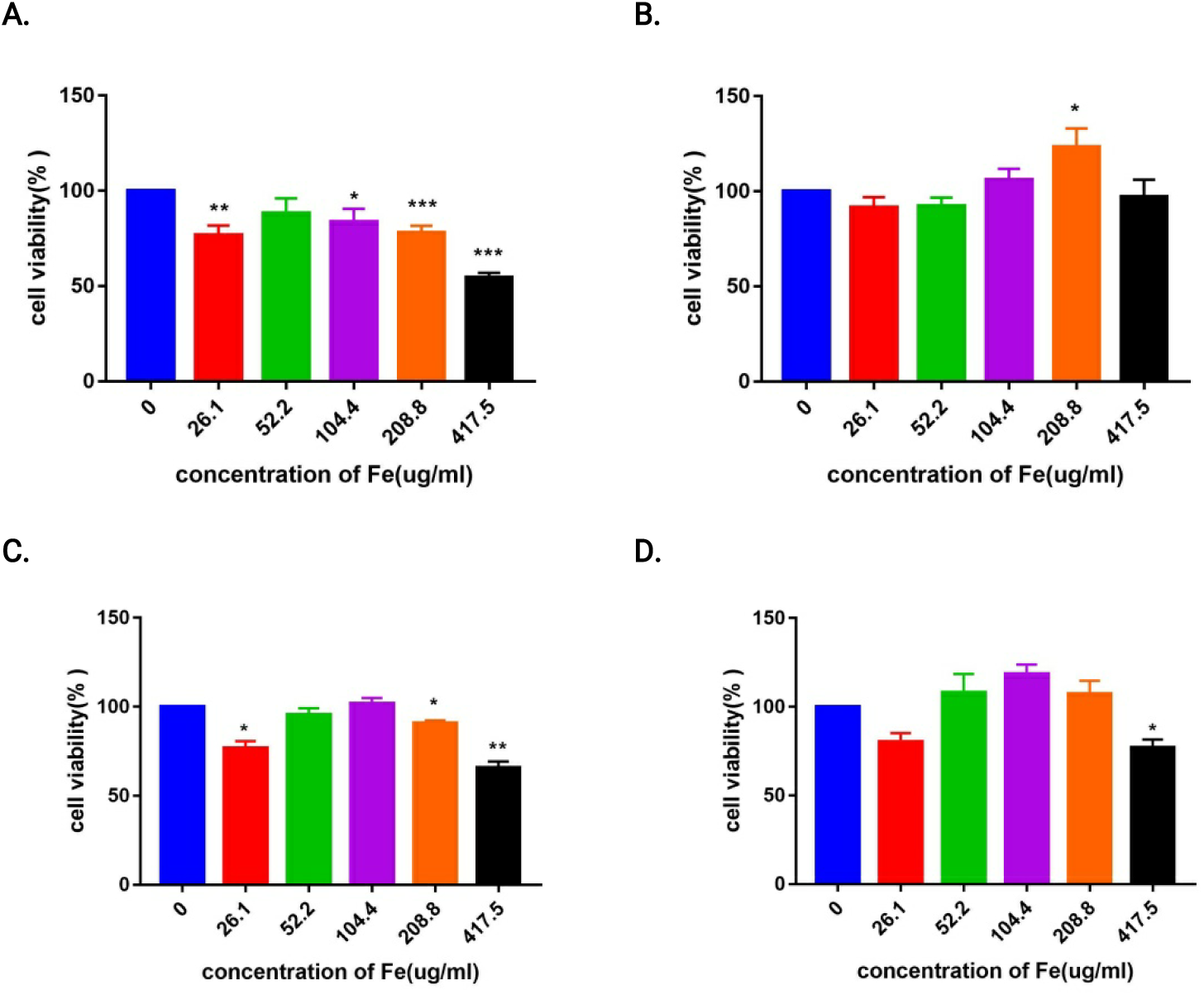
Cell viability was detected by CCK-8 assay after treated for 5 days at different concentrations, data were normalized to control group (no SPIO exposure). A. No RMF; B. under RMF at 2.5Hz; C. under RMF at 5Hz; D. under RMF at 10Hz. All bars represent mean ± SE. * p < 0.05 vs. control group, ** p < 0.01 vs. control group, *** p < 0.001 vs. control group.

### 2. Established the CUMS model successfully and verified the antidepressent effects of SPIO+10Hz RMF with SPT and FST

#### CUMS modeling

After 35 days of stimulations, we performed SPT and FST again and found that the sucrose preference and immobility were significantly changed before and after modeling (Figure 3 A, B). CUMS insensitive mice with change value of two tests less than 10% compared to the baseline were excluded. The modeling success rate is about 60-70%.

**Figure 3 A. and B.**
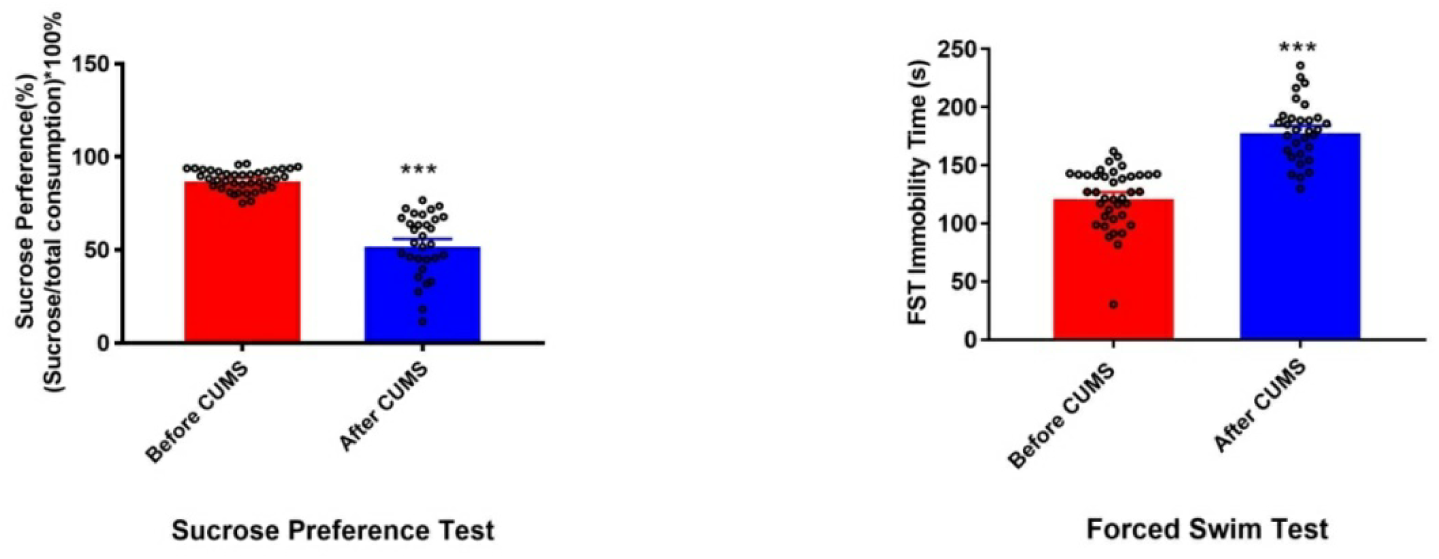
SPT and FST experimental results of mice with CUMS modeling. Values represent the mean ± SE, *** p < 0.001 vs. Wild type.

#### SPT

It is shown that SPIO+10Hz treatment has obvious therapeutic effect on CUMS mice [F(3,30) = 13.545, P < 0.001]. Compared with Wild type mice, there is significant decrease in sucrose preference rate of CUMS mice (n=8, p<0.001 vs. Wild type) and the sucrose preference rate increased markedly after treatment (n=9, p<0.01 vs. CUMS). The increase was not observed in CUMS+SPIO group (n=8, p<0.001 vs. Wild type)(Figure 4 A).

**Figure 4 A. and B.**
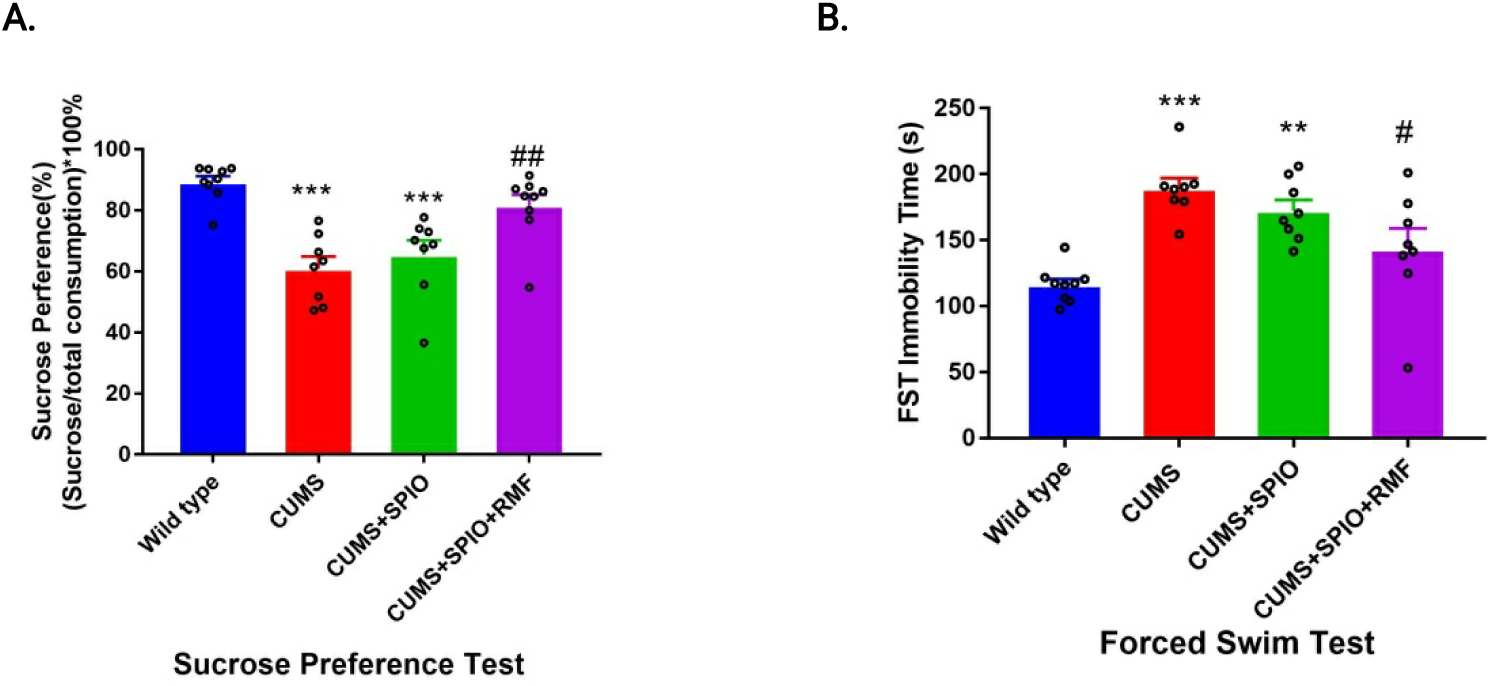
SPT and FST experimental results of mice after treatment. Values represent the mean ± SE. ** p< 0.01 vs. Wild type, *** p < 0.001 vs. Wild type. # p < 0.05 vs. CUMS, ## p<0.01 vs. CUMS.

#### FST

Data from the FST revealed a significant main effect for SPIO+10Hz treatment [F(3,29) = 11.495, P < 0.001]. ANOVA analysis indicated that the immobile time increased significantly after modeling (n=8, P<0.001 vs. Wild type), and the intervention of SPIO + RMF reduced the immobile time (n = 8, P < 0.05 vs. CUMS). However, the CUMS+SPIO group showed no improvement (n=8, P<0.01 vs. Wild type) (Figure 4 B).

### 3. SPIO+10Hz RMF restores the decreased hippocampal BDNF levels induced by CUMS

#### Western Blot

SPIO+10Hz RMF treatment restores the decreased hippocampal BDNF levels induced by CUMS. One-way ANOVA revealed significant effects for treatment [F(3,16)=11.15, P<0.001]. Differences between groups were assessed using Bonferroni Post Hoc Tests. The BDNF level in the hippocampus was significantly down-regulated by CUMS modeling (n=5, P<0.01 vs. Wild type) and the tendency was reversed by SPIO+10 Hz RMF treatment (n=5, P<0.01 vs. CUMS). Merely SPIO failed to up-regulate the protein expression (n=5, P=<0.05 vs. Wild type) (Figure 5 A, B).

**Figure 5.**
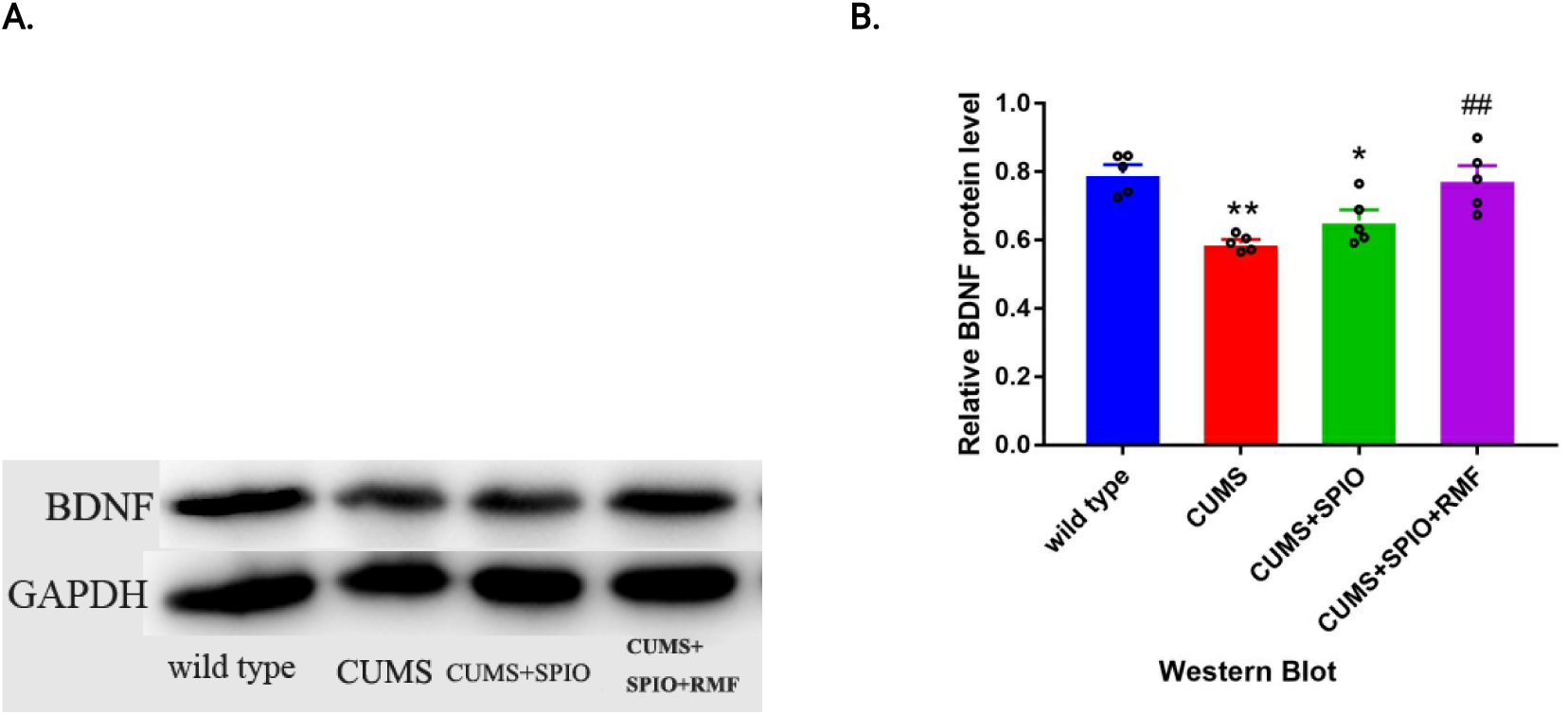
Effects of SPIO+RMF on chronic unpredictable mild stress (CUMS)-induced impairment of hippocampal BDNF. Values represent the mean ± SE. * P<0.05 vs. Wild type, ** p< 0.01 vs. Wild type, ## p<0.01 vs. CUMS.

## Discussion

In short, we proved our hypothesis that micro-nano magnetic particle injected into the hippocampus of mice can produce rapid antidepressive-like effects. During the development of hippocampal neurons, the combinations of SPIO particles with a certain concentration gradient and rotating magnetic field with different frequencies did not display obvious toxic effects. On the other hand, the results from SPT and FST shown that the treatment of 300nl SPIO+10Hz RMF can improve the depressive behavior. What’s more, it is shown that SPIO+10Hz RMF attenuated the impairment of BDNF in hippocampus induced by CUMS. The most interesting thing is that we found the results of FST and BDNF expression in the mice treated with bilateral injection and RMF intervention were not superior to that of the mice treated with unilateral injection.

It is assumed that the regional magnetic field, induced by the magnetization of SPIO particles in brain under the interference of external magnetic field, can enhance the intensity of magnetic stimulation in the hippocampus. Data from other experiments suggest that magnetic stimulation can promote hippocampal neurogenesis and enhance synaptic plasticity [43, 44]. Feng et al. observed that high-frequency rTMS promoted cellular proliferation in the rat hippocampus [45]. Guo et al. demonstrated that rTMS can ameliorate cognitive impairment in PSCI model by enhancing neurogenesis and suppressing apoptosis in the hippocampus [46]. While to our knowledge, there is still no experimental study on direct magnetic stimulation of the hippocampus up to now. Evidence from many studies has revealed that BDNF is involved in the survival and proliferation of new neurons [47, 48] and increasing of BDNF level is closely associated with the depression improvement [32, 36]. We supposed that the SPIO+RMF may take effect by affecting hippocampal neurogenesis or affecting BDNF signaling pathway.

In addition to identifying a novel method for achieving fast acting anti-depressive effects, it provide another new means of magnetic stimulation in arbitrary brain regions. As to magnetic stimulation, TMS is seen as a promising magnetic stimulation therapy for depression factually. However the action site of TMS is usually limited to the cortex, especially left prefrontal cortex [49] due to the attenuation property of a magnetic field. Therefore, SPIO would help to investigate the antidepressant effects of deep brain magnetic stimulation. What’s more, owing to the accurate localization of SPIO in brain regions, the spatial resolution of magnetic stimulation is improved, which would be of great benefit to study the anti-depression mechanism of magnetic stimulation in mice or rats.

The current results show that combination of SPIO in brain and magnetic field outside may provide a unique and fast strategy for depression treatment. However, more experiments with large samples are still necessary to verify its effectiveness. On the other hand, in order to take place of TMS to do research on the mechanism of magnetic stimulation therapy in mice or rats, it is indispensable to verify the biphasic characteristics of this novel treatment in advance. Finally, our findings suggest that SPIO+RMF would be promising in depression treatment and the study on mechanism of magnetic stimulation therapy.

